# Targeted Sequencing Workflows for Comprehensive Drug Resistance Profiling of *Mycobacterium tuberculosis* cultures using Illumina MiSeq and Nanopore MinION: Comparison of analytical and diagnostic performance, turnaround time and cost

**DOI:** 10.1101/760462

**Authors:** Ketema Tafess, Timothy Ting Leung Ng, Hiu Yin Lao, Kenneth Siu Sing Leung, Kingsley King Gee Tam, Rahim Rajwani, Sarah Tsz Yan Tam, Lily Pui Ki Ho, Corey Mang Kiu Chu, Dimitri Gonzalez, Chalom Sayada, Oliver Chiu Kit Ma, Belete Haile Nega, Gobena Ameni, Wing Cheong Yam, Gilman Kit Hang Siu

## Abstract

The emergence of *Mycobacterium tuberculosis* strains with complex drug resistance profiles necessitates a rapid and extensive drug susceptibility test for comprehensive guidance of patient treatment. Here, we developed two targeted-sequencing workflows based on Illumina MiSeq and Nanopore MinION for the prediction of drug resistance in *M. tuberculosis* towards 12 anti-tuberculous agents.

A total of 163 *M. tuberculosis* cultured isolates collected from Hong Kong and Ethiopia were subjected to a multiplex PCR for simultaneous amplification of 19 drug-resistance associated genetic regions. The amplicons were then barcoded and sequenced in parallel on MiSeq and MinION in respective batch sizes of 24 and 12 samples. Both platforms successfully sequenced all samples with average depths of coverage of 1,127× and 1,649× respectively. Utilizing a self-developed Web-based bioinformatics pipeline, Bacteriochek-TB, for variant analysis, we found that the MiSeq and MinION result could achieve 100% agreement if variants with an allele frequency of <40% reported by MinION were excluded. For drug resistance prediction, both workflows achieved an average sensitivity of 94.8% and specificity of 98.0% when compared with phenotypic drug susceptibility test. The turnaround times for the MiSeq and MinION workflows were 38 and 15 hours, facilitating the delivery of treatment guidance at least 17-18 days earlier than pDST respectively. The higher cost per sample on the MinION platform (US$71.56) versus the MiSeq platform (US$67.83) was attributed to differences in batching capabilities.

Our study demonstrated the interchangeability of MiSeq and MinION sequencing workflows for generation of accurate and actionable results for the treatment of tuberculosis.

**Importance:** TB therapy involving different combinations of antibiotics have been introduced to address the issue of drug resistance. However, this practice has led to increasing numbers of *M. tuberculosis* with complex drug resistance profiles. Molecular assays for rapid and comprehensive drug resistance profiling of *M. tuberculosis* are lacking.

Here, we described targeted-sequencing workflows based on Illumina MiSeq and Nanopore MinION for the detection of drug resistance mutations scattered across 19 genetic regions in *M. tuberculosis*. A bioinformatics pipeline was also developed to translate raw datasets into clinician-friendly reports that provide comprehensive genetic information for the prediction of drug resistance towards 12 antibiotics.

This is the first study to evaluate and compare the uses of Illumina and Nanopore platforms for diagnosis of drug-resistant tuberculosis. Remarkably, our diagnostic strategy is compatible with different sequencing platforms that can be applied in diagnostic centres with different levels of throughput and financial support for TB diagnosis.

## Introduction

Tuberculosis (TB) remains an enormous public health challenge. Epidemics of TB are fuelled by the emergence and spread of drug-resistant strains of *Mycobacterium tuberculosis*. Despite the initiation of combination therapies, the bacteria develop complex drug resistance profiles, resulting in multidrug-resistant TB (MDR-TB), extensively drug-resistant TB (XDR-TB) and, more recently, totally drug-resistant TB (TDR-TB) (1).

Culture-based phenotypic drug susceptibility tests (pDSTs) are considered the gold standard for determination of drug resistance in *M. tuberculosis*. However, the long turnaround time hinders the delivery of actionable results for early chemotherapeutic guidance (2–4). Molecular diagnostic methods, such as GenoType MTBDRplus (5, 6) and Xpert MTB/RIF assay (7), were developed for the rapid diagnosis of MDR-TB. Nevertheless, these methods are limited to hotspot mutations and provide only partial data on drug susceptibility (8, 9). Data from the World Health Organisation (WHO) indicate that an estimated 50% of drug-resistant TB cases remain undetected and are not treated appropriately (10). Therefore, rapid, accurate and comprehensive diagnostic tools for the identification of drug-resistant TB is remain urgently needed.

The use of next-generation sequencing (NGS) to detect drug resistance associated mutations and enable comprehensive drug resistance profiling represents a promising approach to clinical care. For high-income countries with low TB burdens, whole-genome sequencing (WGS) is an attractive approach because genomic data can be used for transmission tracing and epidemiological investigations, as well as the determination of drug resistance (11, 12). However, in TB-endemic low-middle income countries (LMICs), sequencing efforts should focus on genetic regions associated with drug resistance to maximise sample batching in a single sequencing run. Amplicon-targeted sequencing is a more appropriate option in this scenario(13).

Currently, Illumina MiSeq is the most widely used platform for clinical applications (8, 11, 12, 14). Despite its accuracy and large data output, the high instrument and maintenance costs may not be affordable by clinical centres intended for individualized care.

The Nanopore MinION, a recently introduced sequencing device, is portable, reasonably priced and lacks maintenance requirements (15). This device requires only a laptop computer for operations and can be set up easily in any laboratory or regional chest clinic. However, the low sequencing accuracy is a well-known disadvantage of this platform (16, 17). A comprehensive evaluation of the applicability and reliability of the MinION platform for the diagnosis of drug-resistant TB remains lacking.

A new diagnostic strategy compatible with different sequencing platforms that can be applied in diagnostic centres with different levels of throughput and financial support is beneficial to global control of drug-resistant TB. To this end, we developed two targeted-sequencing workflows based on Illumina MiSeq and Nanopore MinION, respectively, for the detection of drug resistance mutations scattered across 19 genetic regions in *M. tuberculosis*. A Web-based bioinformatics pipeline was also developed to translate raw sequencing datasets into clinician-friendly reports that provide comprehensive genetic information for the prediction of drug resistance towards 12 anti-tuberculous agents. The two platforms were assessed and compared in terms of the sequencing performance, agreement in variant calling, reagent costs and time to reporting. The diagnostic performance of the sequencing results was also evaluated against pDST. To the best of our knowledge, this is the first study to evaluate and compare the uses of Illumina and Nanopore platforms for drug resistance profiling of *M. tuberculosis* using targeted-sequencing approach.

## Materials and Methods

### *M. tuberculosis* Clinical Isolates

The study included 163 *M. tuberculosis* clinical isolates collected from Ethiopia and Hong Kong. Seventy isolates were collected from patients at the TB clinic of Asella Hospital in Ethiopia between January and September 2015. The remaining 90 isolates were collected at TB laboratory of Hong Kong West Cluster of Hospital Authority, which specialises in chest medicine and TB, between April 2003 and January 2017. All isolates were stored at −80°C and revived using the Mycobacterial Growth Indicator Tube 960 (MGIT 960) system at 37°C for 14 days prior to analysis.

#### DNA Isolation and purification

Fourteen-day *M. tuberculosis* cultures in MGIT broth were heat-killed at 80°C for 60 min. Crude DNA extracts were prepared using the alkaline lysis method, as previously described (18). DNA extracts from Ethiopia were shipped to Hong Kong for subsequent processing. All DNA extracts were then purified using 1.8× AMPure XP beads (Beckman Coulter, US).

#### Drug Resistance Mutation Panel

Our targeted sequencing platforms focused on 19 genetic regions which were associated with phenotypic resistance to 12 anti-TB agents, namely isoniazid (INH), rifampicin (RIF), ethambutol (EMB), pyrazinamide (PZA), ofloxacin (OFX), moxifloxacin (MOX), kanamycin (KAN), amikacin (AMK), capreomycin (CAP), streptomycin (STR), linezolid (LZD) and bedaquiline (BDQ).

A drug resistance mutation panel was constructed and incorporated into our sequencing analysis pipeline, Bacteriochek-TB, for automatic reporting of drug resistance mutations.

Initially, we created the panel based on the three published databases (TBDReaMDB, MUBII-TB-DB, RESEQTB.ORG) because these constitute a comprehensive resource for drug resistance mutations in *M. tuberculosis* and include all studies published from January 1966 to March 2013 (19–21). However, the information in these databases is limited to first-line drugs and several second-line drugs. Therefore, we also searched for additional publications that reported mutations associated with resistance to CAP, LZD and BDQ, as well as all literature related to resistance mutations in *M. tuberculosis* since the release of the latest databases (March 2013–January 2019). All publications were reviewed carefully to ensure the consistency of information regarding the mutated nucleotides and amino acids (22–43). A flowchart of the process of selecting mutations for our panel is presented in Supplementary Figure 1. Eventually, 267 genetic mutations scattered across 19 loci were included in our panel (Table 1 and **Supplementary Table 1**).

**Table 1.**
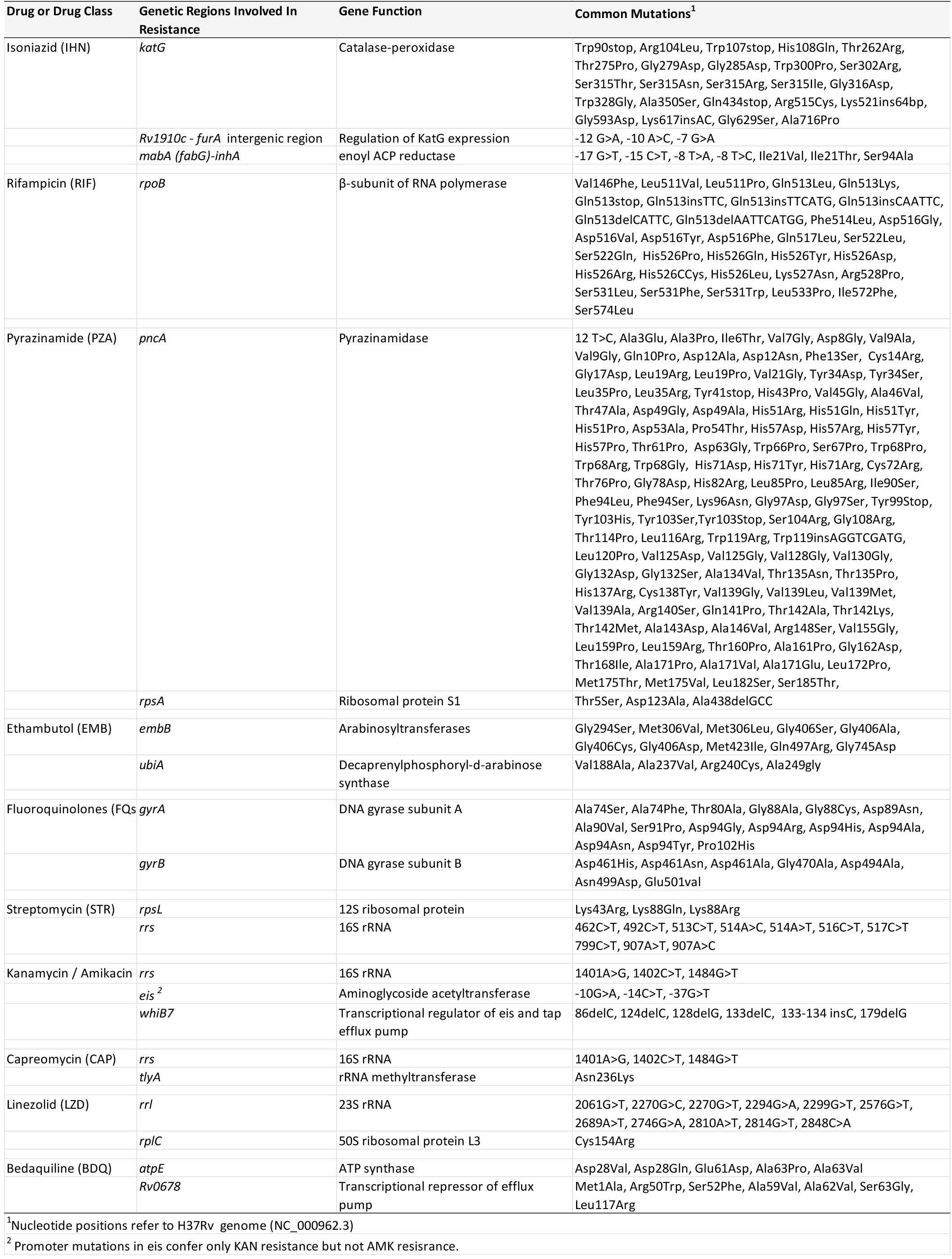
The target loci and mutations included in our drug resistance mutation panel

#### Multiplex PCR

A multiplex PCR was developed for simultaneous amplification of 19 genetic loci. Primers for each targeted locus were designed to amplify amplicon sizes ranging from a minimum of 459 bp (*atpE*) to a maximum of 1601 bp (*rpsA*). Separate sets of primers were designed for the nanopore MinION and Illumina MiSeq sequencing workflows. Primers for MinION sequencing contained a universal tail at the 5’ end to facilitate barcoding PCR and downstream library preparation (Fig.1), while primers for Illumina MiSeq contained only target-specific regions. The sizes of the targeted loci varied by an average of 63 bp to ensure that distinct bands would be visible during electrophoresis (**Supplementary Fig.2**). During multiplex PCR, the target loci were amplified in a total reaction volume of 50 µl, which included 1X KAPA HiFI Hotstart Ready Mix (Roche Sequencing and Life Science, USA), 0.5 µM of each primer (final concentration), 12% GC enhancer (New England BioLabs, USA), and 2 µl of purified DNA extract. Amplification was conducted with the following settings: an initial activation at 95°C for 4 min, 35 cycles of denaturation at 95°C for 30 s, annealing at 63°C for 90 s and extension at 72°C for 90 s and a final extension step at 72°C for 10 min.

**Figure 1:**
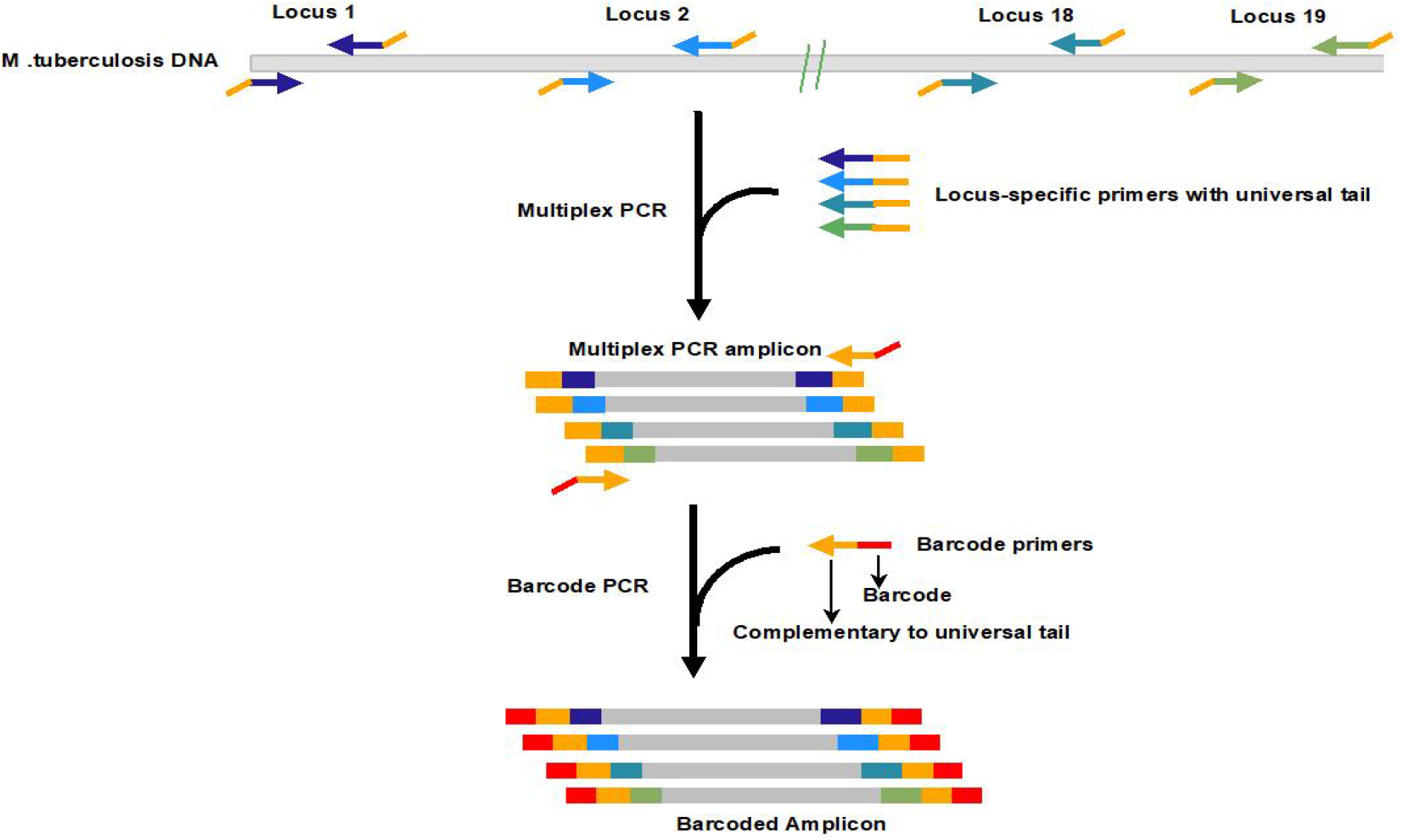
Schematic diagram illustrating multiplex and barcoding PCR amplification for Nanopore MinION platform. Genomic DNA was first amplified with primers containing a universal tail. The 19-plex amplicons were then indexed with barcodes specific to the universal tail that allows the pooling of 12 samples.

#### Illumina MiSeq Sequencing

DNA libraries were prepared and barcoded using the QIAGEN® QIAseq FX DNA Library Kit (Qiagen, USA) according to the manufacturer’s instructions. The quantity and quality of the DNA library were checked using Qubit fluorimeter and Agilent Bioanalyzer 2100, respectively. The final library input for cluster formation was determined using the QIAseq™ Library Quant Assay Kit (Qiagen). The library was normalised, pooled, denatured, spiked with PhiX Control V3 (5%) to ensure base diversity in the amplicons and sequenced using MS-103-1003 MiSeq Reagent Nano Kit version 2 (500 cycles) sequencing chemistry (2 × 250bp) (Illumina). In our clinical settings, the average *M. tuberculosis* positive culture rate was approximately 20–30 per week. Therefore, the workflow was optimised for a batch run of 24 barcoded libraries.

#### Nanopore MinION Sequencing

All samples were analysed using ligation-based 1D sequencing approach (SQK_LSK108). In brief, multiplex PCR amplicons were barcoded using the PCR Barcoding Expansion 1-12 kit (EXP-PBC001; Oxford Nanopore Technologies, UK) according to the manufacturer’s instructions. Twelve barcoded PCR amplicons per batch were normalised and pooled to yield a final DNA quantity of 1 µg. End-repair and dA-tailing were performed using the NEBNext® Ultra™ II End Repair/dA-Tailing Module (New England Biolabs, USA), followed by bead purification and elution with nuclease-free water. A minimum pooled library amount of 700 ng was adapter-ligated using Blunt/TA Ligation Master Mix (New England BioLabs), followed by purification using 0.4× AMPure XP beads. The purified library was then loaded into a R9.4.1 flow cell (FLO-MIN106) and sequenced for 6 hours.

#### Sequencing analysis

The datasets generated from both platforms were analysed using BacterioChek-TB, a CE-IVD marked (LU/CA01/IVD/69) Web-based bioinformatics developed by our team. The software is compatible with targeted or whole-genome sequencing data in the FASTQ format. After inputting the FASTQ file to the software via drag-and-drop, the sequences were mapped against *M. tuberculosis* H37Rv (GenBank accession no. NC_000962.3) using the Burrows-Wheeler Aligner (BWA), version 0.7.15. Expert System, an in-house algorithm, was used to call nucleotide and amino-acid variants from a pileup of aligned read bases. Low-quality variants were removed using a system of adjustable filters, including (i) a noisy mutation filter, which removes variants called with a frequency lower than the limit of detection (3% by default) of the sequencing platform, and (ii) an allele frequency filtering, which excludes mutations with an allele frequency below a defined threshold (10% by default) from the report.

The variants which passed the filters were then mapped using the mutation panel to identify the presence of drug resistance mutations.

After the analysis, the software automatically generated two reports. The first, a one-page clinical report, included the presence of drug resistance mutations with their allele frequencies and drug resistance interpretations (**Supplementary Fig.3**). The second, a full report, included all the above information plus other details such as the read coverage of each target region and the frequencies of insertions, deletions and substitutions at each position across the targeted regions, regardless of their relevance to drug resistance (**Supplementary Fig.3**).

#### Variant calling agreement between Illumina MiSeq and Nanopore MinION

The default allele frequency threshold defined by BacterioChek-TB for variant calling were based on the sequencing accuracy of the Illumina platforms. These settings were not suitable for Nanopore sequencing, which has a higher basecalling error. To determine the optimal threshold for nanopore sequencing, the MinION datasets of a random subset of 20 isolates in our collection were analysed repeatedly using BacterioChek-TB with various allele frequency threshold levels ranging from 10% to 50%. The variant calling results at each threshold level were compared with those generated by the Illumina MiSeq platform at default threshold (10%). The threshold settings resulting in the highest level of agreement between MiSeq and MinION were then used for the analysis of the remaining 143 isolates.

#### Analytical sensitivity of MiSeq and MinION platforms to detect mutations in mixed populations of drug-susceptible and drug-resistant isolate

Genomic DNA were extracted from a wildtype clinical isolate (ETH_125) and a XDR-TB clinical isolate (WC-33), which harboured drug resistance-conferring mutations in *katG*, *rpoB, rrs, embB, gyrA*, and *rpsL*. DNA concentration was measured and standardised using the Qubit dsDNA HS assay. WC_33 DNA was then diluted sequentially with ETH_125 DNA such that the mutant allele proportion reduced from 100% to 0%. Three to four replicates of each dilution were prepared for sequencing. The average frequencies of mutant alleles in each dilution was determined using the two sequencing platforms.

#### Phenotypic Drug Susceptibility Tests

pDSTs were conducted using two methods. For first-line agents, namely RIF (1 µg/ml), INH (0.1 µg/ml and 0.4 µg/ml), EMB (5 µg/ml), STR (1 µg/ml) and PZA (100 µg/ml), the pDST was performed using a liquid-based BACTEC MGIT 960 SIRE kit and BACTEC MGIT 960 PZA kit (44, 45). For second-line agents, including OFX (2 µg/ml), MOX (0.5 µg/ml), AMK (4 µg/ml), KAN (5 µg/ml) and CAP (10µg/ml), the agar proportion method was used to determine drug susceptibility patterns according to the guideline of the Clinical and Laboratory Standards Institute (CLSI). All of the *M. tuberculosis* cultures from Hong Kong (n=93) were subjected to the full panel pDST, whereas the isolates from the Ethiopia were only subjected to MGIT 960 SIRE tests. Unfortunately, pDSTs for LZD and BDQ were not performed in this study because of the lack of a standardised pDST protocol and the unavailability of these drugs in the study regions.

### Diagnostic Performance Evaluation

The diagnostic sensitivities and specificities for each type of drug resistance in the BacterioChek-TB reports based on MiSeq and MinION data were determined using standard methods and compared to the gold-standard pDST results. The adjusted Wald method and free online software (available from http://www.measuringusability.com/wald.htm) were used to calculate the 95% confidence intervals (95% CI).

### Turnaround Time and Cost Assessment

The sample-to-report times were determined for the MiSeq and MinION methods. Time zero was defined as the time when the MGIT cultures were subjected to DNA extraction, while the time-to-report referred to the time when BacterioChek-TB generated a genotypic DST report. The turnaround times of the sequencing methods were compared with that of pDST.

All the cost analyses were based on Hong Kong list prices as of July 2019. For the MiSeq workflow, the average cost per sample for library preparation, quality control and sequencing was calculated based on a batch of 24 samples per run. For MinION, the respective cost per sample was calculated based on a batch of 12 samples per run.

### Statistical analysis

The statistical analysis was performed using GraphPad Prism, version 8.1 (GraphPad Inc., USA). Descriptive results, such as the total number of reads per run, number of reads per sample, read quality and depth of coverage per genetic region obtained by MiSeq and Nanopore sequencing, are expressed as average values. Continuous variables were compared using the independent t-test or Kruskal–Wallis test and one-way ANOVA followed by Dunn’s post hoc test. Variables with p values ≤0.05 were considered statistically significant.

## Results

### Quality, Yield and Depth of Coverage of Sequencing Reads Generated by the Illumina MiSeq and Nanopore MinION Platforms

For Illumina MiSeq, 163 samples were divided into seven batches of 24 libraries. More than 90% of the bases received scores >Q30 for the paired-end reads. Each sample yielded an average read number of 106,050 (**Supplementary Table 2**). For Nanopore MinION, the samples were sequenced in 14 batches of 12 libraries. The average quality score was 10.1 per run, which corresponds to an approximate accuracy of 90%. Average reads of 57,766 per sample were obtained after a 6-hour sequencing run (**Supplementary Table 3**).

Both MiSeq and MinION successfully sequenced all 19 targeted regions in the 163 samples, with average depths of coverage of 1,127× and 1,649× respectively. However, we observed that the depth of coverage decreased as the amplicon size increased. For amplicons smaller than 1,000 bp, the average depths of coverage obtained using MiSeq and MinION were 1,408× and 2,386×, respectively, which were significantly higher than the values (i.e. 815× and 830×) obtained from amplicons ≥1,000 bp [t=16.71; P=0.0001 and t=12.67; P=0.0001, respectively] (Fig.2). A strong inverse relationship was observed between the amplicon size and depth of the coverage for both platforms [Pearson correlation (r)=-0.82, p=0.01 and (r)=-0.91, p=0.01, respectively].

**Figure 2:**
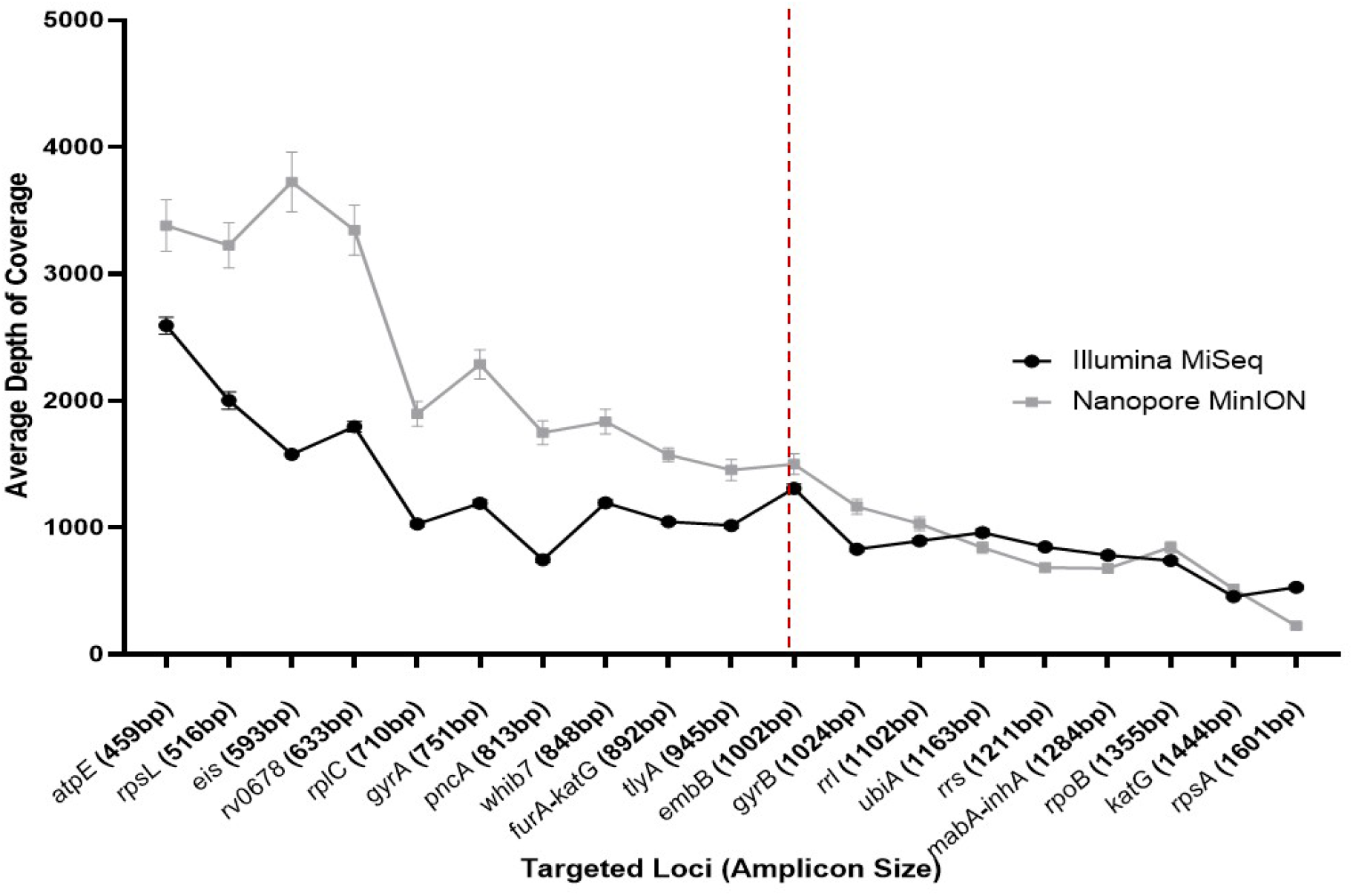
Line graph illustrating the depth of coverage at each locus in MinION and MiSeq sequences (n=163). Error bars: Standard error of mean (SEM); The reference line (broken line) indicates the cut-off point (1,000bp) to categorize the amplicon size as long and short

### Variant Call Percent Agreement between MiSeq and MinION Sequencing

The variant call percent agreements were 1.84%, 16.78%, 56.58%, 100% and 97.67% for MinION at allele frequency threshold of 10%, 20%, 30%, 40% and 50% respectively versus MiSeq using default filter setting (Fig.3). The variant calling results were presented in Supplementary Table 4. The results showed that the MiSeq and MinION result could achieve 100% agreement if variants with an allele frequency of <40% reported by MinION were excluded.

**Figure 3.**
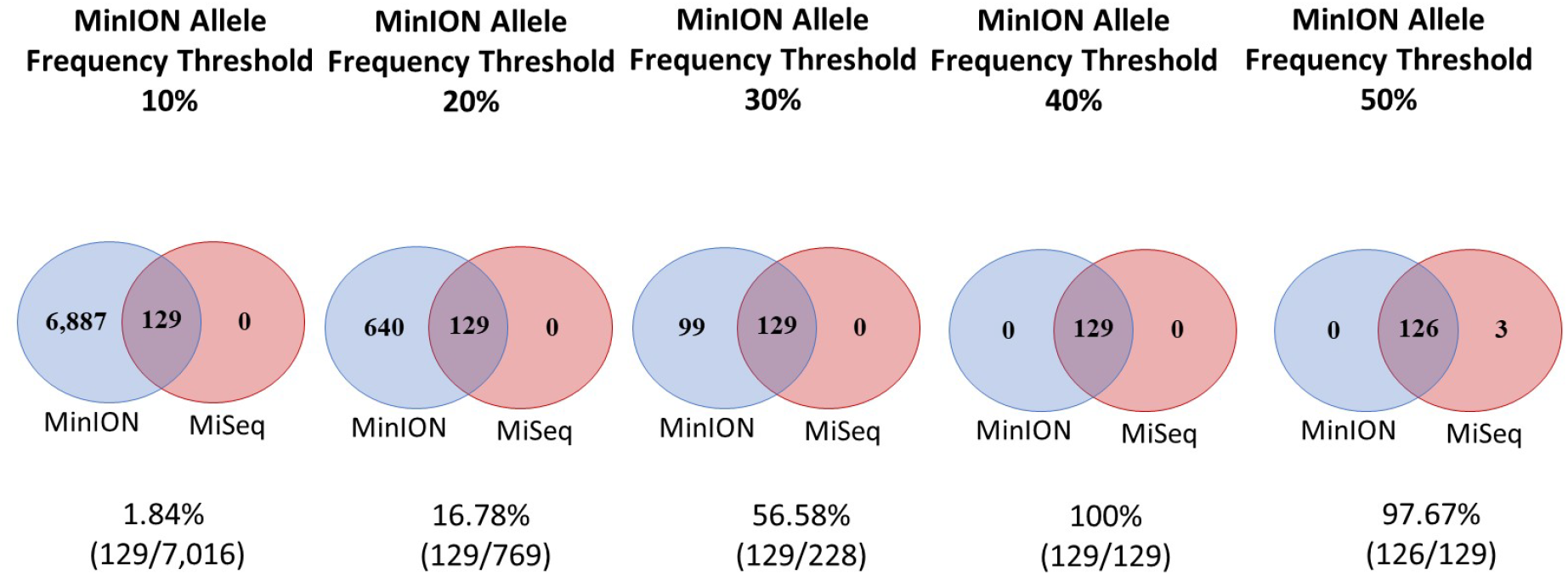
Venn diagrams showing the agreement of variant calling between two sequencing platforms when the allele frequency threshold was adjusted to 10% (a default setting) for Illumina MiSeq and 10-50% for Nanopore MinION. The sum of the number in a circle indicates the total number of variants, regardless of their relatedness with drug resistance, reported by the respective sequencing platform for a subset of 20 *M. tuberculosis* cultures.

### Analytical sensitivity of MiSeq and MinION platforms to detect mutations in mixed populations of drug-susceptible and drug-resistant isolate

Mixtures of genomic DNA extracted from a pan-susceptible isolate and an XDR-TB isolate, which harboured six drug resistance mutations, at different ratios were analysed by MiSeq alongside MinION. Notably, the numbers of reads carrying the mutations increased proportionally with an increasing mutant/wild type ratio (Fig.4). For MiSeq, a drug-resistant mutant DNA proportion of at least 20% was required to exceed the allele frequency threshold of 10% for all six regions. For MinION, a drug-resistant subpopulation comprising at least 50% of the sample mixture was required to achieve an allele frequency of 40%.

**Figure 4.**
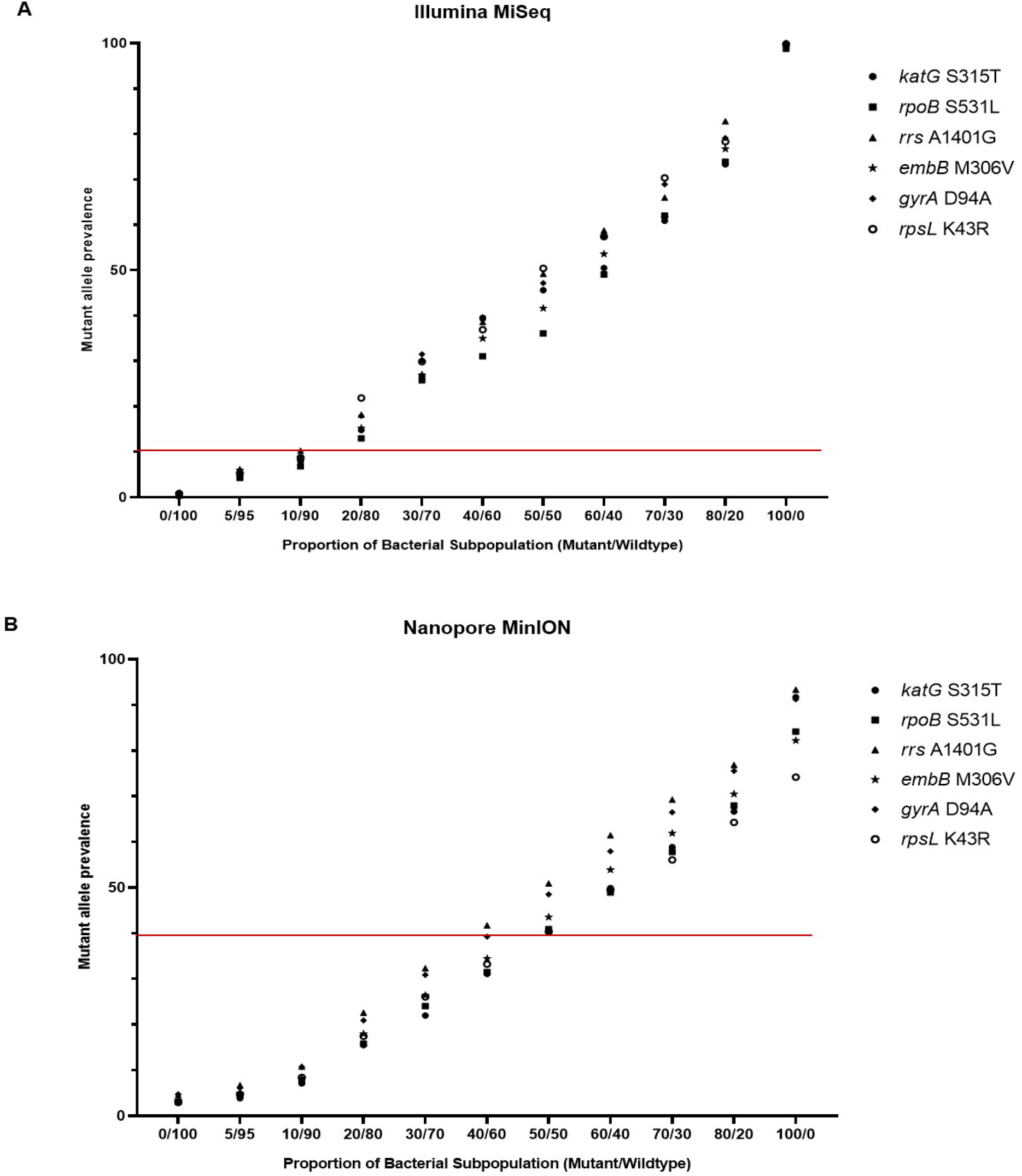
The frequencies of mutant alleles identified by (A) Illumina MiSeq and (B) Nanopore MinION sequencing platforms in a mixed population of wildtype and mutant at different ratios. The mutant strain, WC-33, were found to harbour 6 multiple resistance-conferring mutations (*katG* Ser315Thr, G > C genome position (P) 2155168; *rpoB* Ser531Leu, C > T P 761155; *rrs* A1401G, A > G P 1473246; *embB* Met306Val, A > G P 4247429; *gyrA* Asp94Ala, A > C P 7582; *rpsL* Lys43Arg, A > G P 781687). ETH_125 represent the wildtype strain, which carried pure wildtype alleles at the respective positions. The red line indicates the respective allele frequency thresholds for MiSeq and MinION to report a variant by Bacteriochek-TB. The allele proportion was calculated as the total number of specific allele per total number aligned reads at that specific position for all the replicates.

### Phenotypic Drug Susceptibility Patterns of *M. tuberculosis* Cultures

All 163 *M. tuberculosis* isolates were subjected to the MGIT 960 SIRE test. Of them, 65 (39.9%) isolates were pan-susceptible, 98 (60.1%) were resistant to at least one first-line drug and 60 (36.8%) were categorised as MDR-TB. The resistance rates to individual drugs are listed in Table 2.

**Table 2:**
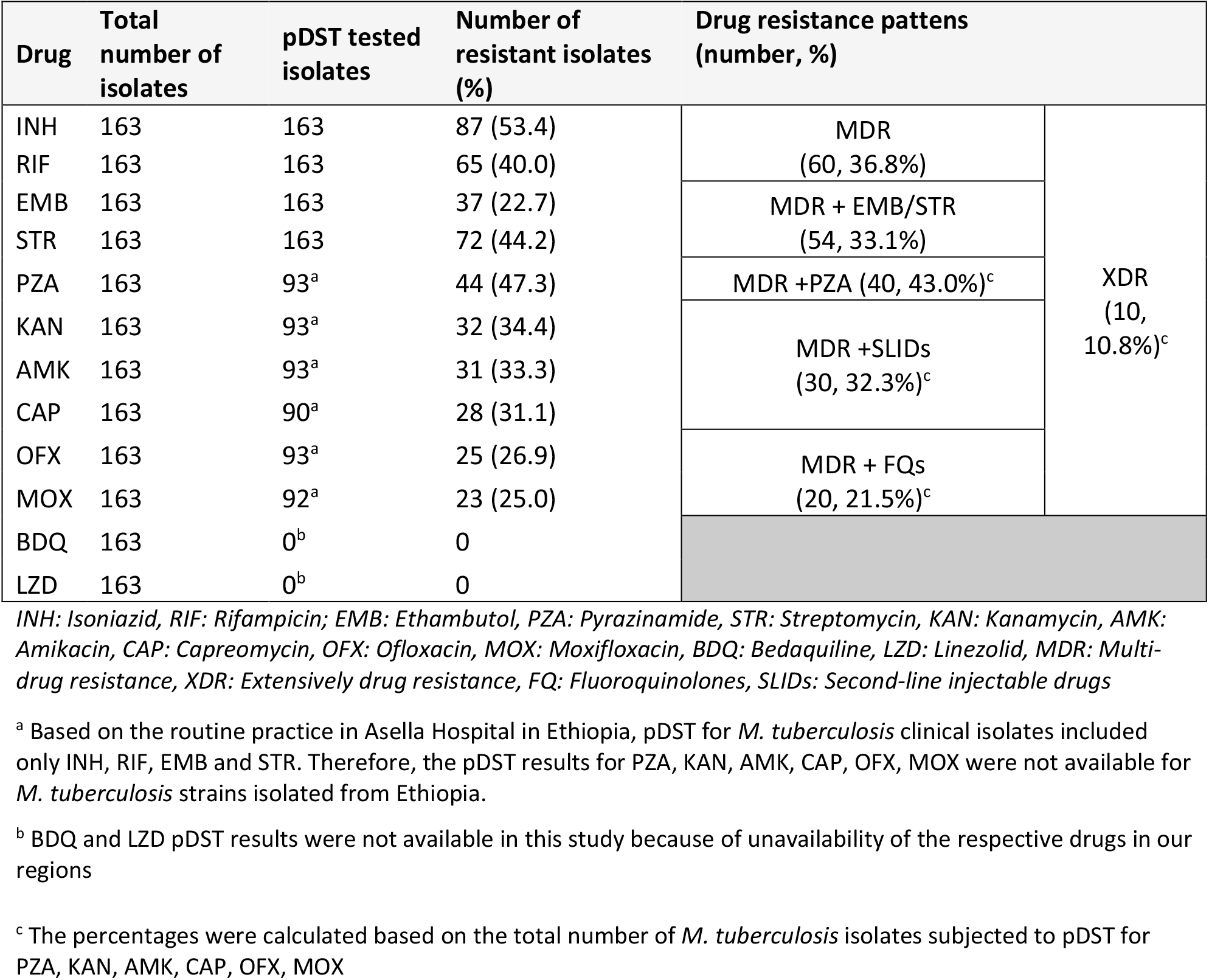
Phenotypic drug resistance profiles of 163 *M. tuberculosis* clinical isolates

Ninety-three *M. tuberculosis* isolates collected from Hong Kong were further tested for susceptibilities toward PZA, OFX, MOX, KAN, AMK and CAP. Of them, 30 (32.3%) were found to be MDR plus resistant to any SLIDs, 20 (21.5%) were MDR plus resistant to FQ and 10 (10.8%) were categorised as XDR-TB.

Unfortunately, pDST data for LZD and BDQ could not be determined in this study because these antibiotics were unavailable at the study sites (Table 2).

### Diagnostic Performance of MiSeq and MinION Sequencing Workflows for Predicting Drug Susceptibility Patterns

By applying the respective allele frequency thresholds, the two sequencing platforms deduced identical genotypic patterns for all 163 isolates. Using pDST as the reference method, the average sensitivity and specificity of the sequencing-based methods for the detection of drug resistance in *M. tuberculosis* cultures were 94.8% (95% CI: 92.3–96.6) and 98.0% (95% CI: 96.8–98.8), respectively. Both sequencing platforms correctly identified all MDR-TB (n=60) and XDR-TB isolates (n=10) in our collection.

Regarding the performance for prediction of resistance towards individual drugs, the diagnostic sensitivities and specificities were presented in Table 3.

**Table 3:** The diagnostic performance of the MiSeq and MinION sequencing workflows

| **Drug** | **pDST tested isolates** | **Number of drug resistant isolates as confirmed by pDST**<br>**(% of the tested isolates)** | **Number of drug susceptible isolates as confirmed by pDST**<br>**(% of the tested isolates)** | **Number of drug resistant isolates as inferred by sequencing-based method**<br>**(% of the tested isolates)** | **Number of drug susceptible isolates as inferred by sequencing-based method**<br>**(% of the tested isolates)** | **Sensitivity %**<br>**(95% CI)** | **Specificity %**<br>**(95%CI)** |
| --- | --- | --- | --- | --- | --- | --- | --- |
| INH | 163 | 87 (53.4) | 76 (46.6) | 80 (49.1) | 76 (46.6) | 92.0 (84.1-96.3) | 100.0 (95.9-100.0) |
| RIF | 163 | 65 (40.0) | 98 (60.0) | 65 (40.0) | 97 (59.5) | 100.0 (95.2-100.0) | 99.0 (93.9-99.9) |
| EMB | 163 | 37 (22.7) | 126 (77.3) | 32 (19.6) | 123 (75.5) | 86.5 (71.6-94.6) | 97.6 (92.9-99.5) |
| STR | 163 | 72 (44.2) | 91 (55.8) | 67 (41.1) | 91 (55.8) | 93.1 (84.4-97.3) | 100.0 (96.5-100.0) |
| PZA | 93 | 44 (47.3) | 49 (52.7) | 42 (45.2) | 49 (52.7) | 95.5 (84.0-99.6) | 100.0 (93.7-100.0) |
| KAN | 93 | 32 (34.4) | 61 (65.6) | 31 (33.3) | 61 (65.6) | 96.9 (82.9-99.9) | 100.0 (94.9-100.0) |
| AMK | 93 | 31 (33.3) | 62 (66.7) | 31 (33.3) | 62 (66.7) | 100.0 (90.4-100.0) | 100.0 (94.9-100.0) |
| CAP | 90 | 28 (31.1) | 62 (68.9) | 28 (31.1) | 60 (66.7) | 100.0 (89.5-100.0) | 96.8 (88.3-99.8) |
| OFX | 93 | 25 (26.9) | 68 (73.1) | 23 (24.7) | 68 (73.1) | 92.0 (73.9-98.9) | 100.0 (95.4-100.0) |
| MOX | 92 | 23 (25.0) | 69 (75.0) | 21 (22.8) | 68 (73.9) | 91.3 (72.0-98.8) | 98.6 (91.5-99.9) |

INH: Isoniazid, RIF: Rifampicin; EMB: Ethambutol, PZA: Pyrazinamide, STR: Streptomycin, KAN: Kanamycin, AMK: Amikacin, CAP: Capreomycin, OFX: Ofloxacin, MOX: Moxifloxacin

Missense mutations in codon 315 of *katG* and a C-15T nucleotide change in the *mabA*-*inhA* promoter region were the most prevalent mutations, accounting for 39.1% and 47.1% of INH resistance. Majority (75.4%) of RIF-resistant isolates harboured a missense mutation in codon 531 of *rpoB* (Table 4). Mutations in *embB* and *rpsL* that resulted in the amino acid changes Met306Val and Lys43Arg, respectively, accounted for 75.7% and 68.1% of EMB-and STR-resistant isolates, respectively. Mutations associated with PZA resistance were scattered across *pncA*, in which mutation encoding the amino acid change Gly162Asp was predominant (Table 4).

**Table 4:**
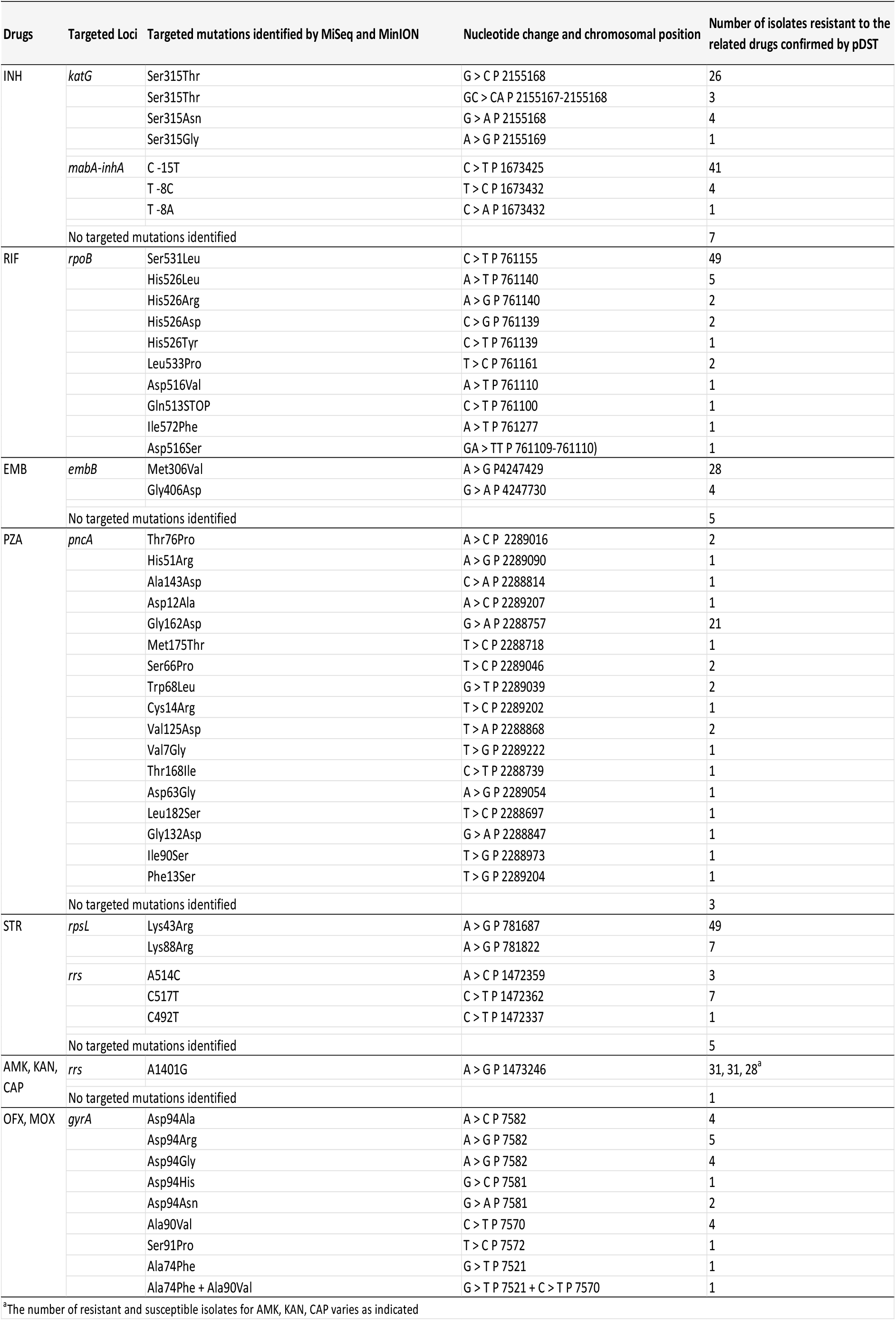
Targeted mutations identified by MiSeq and MinION sequencing workflows for *M. tuberculosis* isolates that were phenotypically resistant to the related drugs

For second-line drugs, the majority (64%) of FQ-resistant isolates harboured missense mutations in codon 94 of *gyrA*, whereas a nucleotide substitution A1401G in *rrs* accounted for nearly all resistance to SLIDs. Detailed phenotypic and genotypic drug susceptibility information of each *M. tuberculosis* isolate were presented in Supplementary Table 5.

### Discordance between pDST and Sequencing-Based Workflows

Discordant genotypic and phenotypic resistance data were observed for some isolates (Table 5). Seven isolates which were phenotypically resistant to INH were inferred as susceptible by both sequencing workflows. Four of these isolates were found to harbour *katG* mutations outside our target panel and three of them did not have any variants identified in related targeted regions. Moreover, an isolate harboured a high-confidence mutation at codon 526 of *rpoB* was found to be phenotypically susceptible to RIF.

**Table 5:** Discordance noted in genotypic and phenotypic tests

| Table 5: Discordance noted in genotypic and phenotypic tests |  |  |  |  |  |  |
| --- | --- | --- | --- | --- | --- | --- |
| Drugs | Targeted Loci | Loci containing the mutations | Mutations | Number of Isolate | Genotypic drug susceptibility inferred by Sequencing-based method | Phenotypic DST Results |
| INH | *katG* , *Rv1910c - furA intergenic region* , *mabA-inhA* promoter | *katG* | Gln461STOP <sup>a</sup> | 1 | Susceptible | Resistant |
|  |  | *katG* | Pro232Leu <sup>a</sup> | 1 | Susceptible | Resistant |
|  |  | *katG* | AAA554AAAA(INSIns) <sup>a</sup> | 1 | Susceptible | Resistant |
|  |  | *katG* | Phe657Ser <sup>a</sup> | 1 | Susceptible | Resistant |
|  |  | No mutations found in all targeted loci |  | 3 | Susceptible | Resistant |
| RIF | *rpoB* | *rpoB* | His526Leu | 1 | Resistant | Susceptible |
| EMB | *embB* , *ubiA* | *embB* | Asp354Ala <sup>a</sup> | 2 | Susceptible | Resistant |
|  |  | *embB* | Glu378Ala <sup>a</sup> | 2 | Susceptible | Resistant |
|  |  | *ubiA* | Glu149Asp <sup>a</sup> |  |  |  |
|  |  | *embB* | Met306Val <sup>a</sup> | 3 | Resistant | Susceptible |
|  |  | No mutations found in all targeted loci |  | 1 | Susceptible | Resistant |
| PZA | *pncA* | *pncA* | 128 GTC > G_C (T_Del P 2288859) <sup>a</sup> | 1 | Susceptible | Resistant |
|  |  | No mutations found in all targeted loci |  | 1 | Susceptible | Resistant |
| STR | *rpsL* , *rrs* | No mutations found in all targeted loci |  | 5 | Susceptible | Resistant |
| CAP | *rrs* , *tlyA* | *rrs* | A1401G | 2 | Resistant | Susceptible |
| KAN | *rrs* , *eis* , *whiB7* | No mutations found in all targeted loci |  | 1 | Susceptible | Resistant |
| FQ | *gyrA* , *gyrB* | *gyrA* | Asp94Arg | 1 | Resistant | Susceptible <sup>b</sup> |
|  |  | *gyrA* | Val235Val <sup>a</sup> | 1 | Susceptible | Resistant |
|  |  | *gyrB* | Ser447Phe <sup>a</sup> | 1 | Susceptible | Resistant |

<sup>a</sup> Out-of-panel (non-targeted) mutations identified in the targeted loci in *M. tuberculosis* isolates with discordant genotypic and phenotypic DST results. <sup>b</sup> The isolate was phenotypically resistant to OFX but susceptible to MOX.

Sequencing-based workflows also failed to identify two (4.5%) PZA-resistant isolates. One isolate harboured a nucleotide deletion in *pncA*, whereas the other did not harbour any mutations in *pncA* or *rpsA*. Both the MiSeq and MinION workflows missed five STR-and EMB-resistant isolates. No known drug resistance mutations were detected in *rrs* and *rpsL* for any of the five STR-resistant isolates, whereas out-of-panel *embB* mutations were detected in four of the EMB-resistant isolates Moreover, three isolates harbouring a high-confidence mutation, *embB* Met306Val, were identified as phenotypically susceptible to EMB.

Two FQ-resistant isolates harboured out-of-panel mutations in *gyrA* and *gyrB*. One isolate carrying a targeted mutation in *gyrA*, which caused the amino acid change Asp94Arg, was resistant to OFX but susceptible to MOX.

Only one (3.2%) SLIDs-resistant isolate was not detected by the sequencing tests. Although this isolate was phenotypically resistant to KAN, no targeted mutations were detected in *rrs*, *eis* or *whiB7.* Two isolates harbouring the high–confidence mutation *rrs* A1401G were shown to be resistant to KAN and AMK but susceptible to CAP.

### Turn-around Times and Cost Assessments of the MiSeq and MinION Sequencing Workflows

The accumulated occupation hours from the DNA extraction from MGIT culture to the generation of a report by BacterioChek-TB were 38 and 15 hours for the MiSeq and MinION workflows, respectively. In a laboratory with daily working hours of 9:00 am to 5:00 pm, genotypic DSTs generated using the MiSeq and MinION workflows could be available in 4 and 3 days, respectively. In other words, the MiSeq and MinION workflows respectively reported drug susceptibility results for first-line agents 9 and 10 days earlier than the 13-day protocol required for MGIT 960 SIRE. A full-panel genotypic DST of 12 anti-TB agents could be delivered for treatment guidance at least 17 and 18 days earlier than the pDST results, respectively (Figure 5).

**Figure 5:**
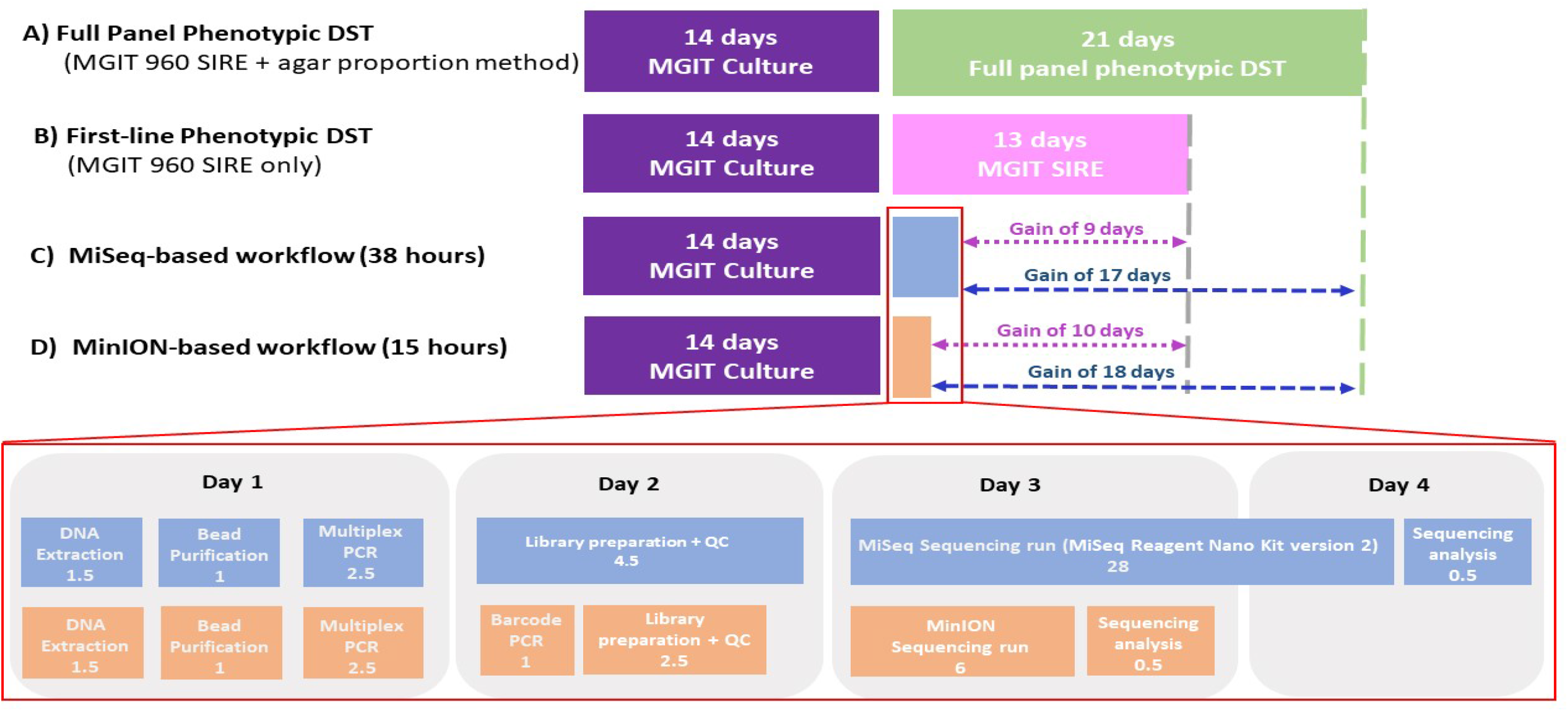
Illustration of the turnaround time of phenotypic drug susceptibility test (pDST) versus our sequence-based assays. Archived *M. tuberculosis* strains were incubated in MGIT broth for 14 days before subjecting to **(A)** a full panel of pDST, including MGIT 960 SIRE (13 days), MGIT 960 PZA (21 days) and agar proportion method for 2^nd^ line drugs (21 days); **(B)** MGIT 960 SIRE only (13 days); **(C)** MiSeq sequencing workflow, which requires about 38 hours – DNA extraction (1.5 hrs), 1.8X bead purification (1 hr), multiplex PCR (2.5hrs), library preparation including fragmentation and adapter ligation, Bioanalyzer and quantitative real-time PCR assessment (4.5 hours), Sequencing run (28 hours) and sequencing analysis (0.5 hrs); **(D)** MinION sequencing workflow, which requires about 15 hours-DNA extraction (1.5 hrs), 1.8X bead purification (1 hr), multiplex PCR (2.5hrs), barcoding PCR (1hr), library preparation plus quality and quantity check (2.5 hrs), sequencing run (6hrs) and sequencing analysis (0.5hrs). Note: The boxes are not to the scale

The running costs per sample (including reagents and consumables) were US$67.83 for MiSeq sequencing (24 samples/run) and US$71.56 for MinION sequencing (12 samples/run) (Table 6).

**Table 6:**
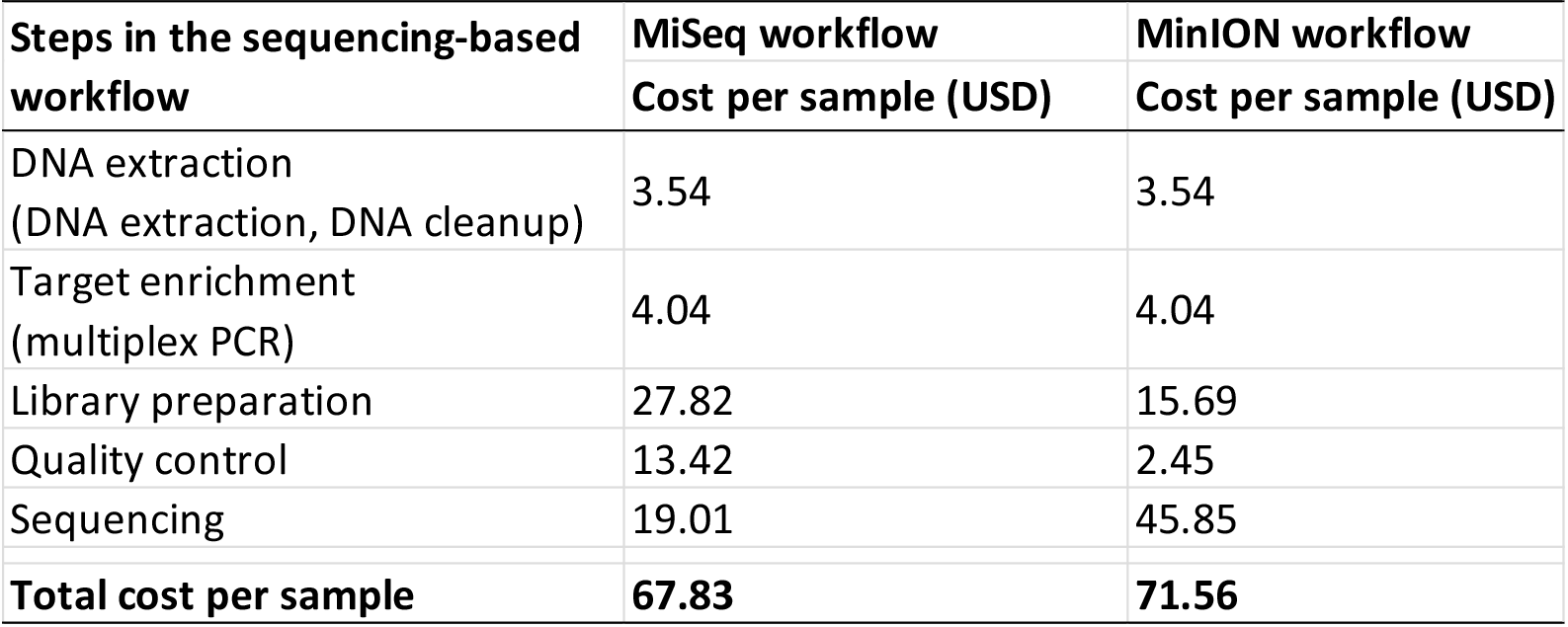
The cost of reagents and consumables per sample of MiSeq and MinION sequencing workflows

|  |  |  |
| --- | --- | --- |
| **Steps in the sequencing-based workflow** | **MiSeq workflow** | **MinION workflow** |
|  | **Cost per sample (USD)** | **Cost per sample (USD)** |
| DNA extraction<br>(DNA extraction, DNA cleanup) | 3.54 | 3.54 |
| Target enrichment<br>(multiplex PCR) | 4.04 | 4.04 |
| Library preparation | 27.82 | 15.69 |
| Quality control | 13.42 | 2.45 |
| Sequencing | 19.01 | 45.85 |
| **Total cost per sample** | **67.83** | **71.56** |

## Discussion

The translation of sequencing-based workflows from research laboratories to clinical settings requires an evaluation of the analytical and diagnostic performance and an assessment of usability in terms of costs and turnaround times. This was the first study to evaluate and compare the uses of Illumina and Nanopore sequencing platforms for the targeted sequencing-based drug resistance profiling of *M. tuberculosis* cultures.

Instead of sequencing the whole genome of *M. tuberculosis* (4.4 Mb), the total region size to be sequenced (i.e., sum of 19 genetic regions) was only 18,346 bp. This facilitates the simultaneous analysis of multiple samples in one flow cell via barcoding on both platforms. In this study, a batch run of 24 samples on MiSeq yielded an average depth of coverage of 1,127× per sample. Nevertheless, the depth of coverage was not consistent across the 19 target regions. The average coverage of long loci could be as low as 455×, while the short loci achieved 2,592×. Likewise, the MinION workflow also yielded an inconsistent depth of coverage across the target loci. While the shortest locus, *atpE*, achieved an average read depth of 3,380×, the longest target, *rpsA*, had an average read depth of only 223×. The low read depths of long loci limited the capacity for scaling up the batch size given that the recommended depth of coverage for each locus in a BacterioChek-TB analysis is 100×. Accordingly, the maximum batch size should be approximately 108 samples per run for MiSeq workflow and 24 samples per run for MinION. The inconsistent coverage could be attributed to amplification bias during multiplex PCR, wherein PCR polymerase preferably amplifies shorter DNA fragments which consequently outnumbered long DNA fragments in the resultant mixture of amplicons.

While the Illumina platform has been deployed widely for clinical purposes (46), the manufacturer of the Nanopore MinION insisted that the current version of the platform was not intended for clinical applications, possibly because of relatively high basecalling error rate. Still, it would be interesting to determine the agreement between the MinION and Illumina platforms in terms of the identification of genetic variants, as this would allow us to explore the potential usefulness of the former for the diagnosis of drug-resistant TB. As the same pipeline was used for the analysis of both datasets, any differences in variant calls should be driven solely by sequencer differences. Initially, we defined an allele frequency threshold of 10% for both MiSeq and MinION and observed a variant call percent agreement of 1.84%. As the reported accuracy of the basecaller (Albacore v.2.3.3) for MinION data was 88%, we expected a good concordance if the threshold level was increased to 20% for MinION. Unfortunately, the agreement only increased to 16.78%. More than 81% of these ‘faked’ variants were identified in *rrs* and *rrl* (**Supplementary Table 4**). The high GC contents and repeating sequences in these regions might have worsened the basecalling accuracy. Perfect agreement between the two platforms was finally achieved at a threshold of 40%, indicating that the allele frequencies of the ‘faked’ variants due to basecalling errors was always <40%. Notably, the percent agreement decreased to 97.67% when the allele frequency threshold was increased further to 50%. Some true variants which were identified at an allele frequency >95.0% by MiSeq were reported by MinION at a prevalence of only 45–48% and were excluded at a threshold level of 50%.

In this study, all 163 *M. tuberculosis* cultures were resuscitated from frozen stocks of pure strains. No heterogenous drug susceptibility was expected. However, in clinical settings, *M. tuberculosis* cultures inoculated from respiratory specimens may contain mixed populations of drug-susceptible and drug-resistant strains. While highly specific variant calling could be achieved by MinION by setting an allele frequency threshold of 40%, the sensitivity of this platform to detect minor variants remained uncertain.

Our results showed that the proportions of drug-resistant subpopulations in heterogeneous samples should be large enough to generate a read depth that exceed the allele frequency threshold in order to report the drug-resistant variants. For instance, the drug-resistant strain should comprise at least 20% of the total population to ensure that all six drug resistance mutations would be detected by the MiSeq workflow at the frequency threshold of 10%. Similarly, for the MinION workflow, the drug-resistant subpopulation had to be increased to 50% of the total to ensure that all drug-resistance mutations were reported at a frequency of ≥40%.

The MiSeq workflow obviously outperformed the MinION workflow in terms of the analytical sensitivity to detect minor drug-resistant variants. Previous studies (13, 47) have demonstrated that MiSeq sequencing can identify ultralow heteroresistance, defined as a level significantly lower than the current threshold. Conversely, the MinION workflow would probably miss resistance caused by low-frequency alleles (<40%). Our finding was similar to that of Ammar *et al*., who reported the detection of variants from heterogeneous samples using MinION only if their frequencies > 34% (48).

One key feature of this study was the inclusion of numerous highly resistant *M. tuberculosis* isolates (including MDR-TB, n=60 and XDR-TB, n=10) with diversified drug resistance mutation patterns. Using pDST as a reference standard, both workflows achieved an average sensitivity of 94.8% and specificity of 98.0%, which were higher than the targets set by the WHO for new molecular assays (90% and 95%, respectively) (49). Most importantly, all MDR-TB and XDR-TB isolates in our collection were successfully identified.

The sequencing-based method also correctly identified drug-resistant isolates harbouring rare drug resistance mutations, such as *rpoB* Ill572Phe, *rrs* C492T and *gyrA* Ala74Phe, which were generally outside the target panels of the existing molecular tests (5–7, 50). In addition, molecular diagnostic tools for genotypic predictions of PZA resistance are lacking. It is because the drug resistance variants are scattered across the *pncA* gene and no hotpot mutations are known. Notably, our sequencing-based methods identified 95.5% (42/44) of the PZA-resistant isolates, which exhibited 17 different mutation patterns.

Discordant genotypic and phenotypic resistance data were observed for some isolates, indicating that the genetic mechanisms underlying drug resistance in *M. tuberculosis* are yet to be fully elucidated. Specifically, a significant number of drug-resistant isolates carrying out-of-panel or novel mutations in the targeted loci were not reported as resistance by either sequencing method. For instance, four INH-resistant isolates were found to harbour novel *katG* variants. The causative roles of these mutations in conferring INH resistance remain to be validated experimentally. They will be included in the panel once their roles have been confirmed. Similar follow-up studies will be performed to validate the causative roles of novel variants in *embB*, *pncA*, *gyrA* and *gyrB* identified in this study. Consequently, the drug resistance mutation panel will be expanded continuously to improve the diagnostic performance.

Moreover, some drug-resistant isolates did not harbour any mutations in the targeted genetic regions. For instance, 6.9% (5/72) of STR-resistant isolates harboured no known mutations across *rrs* and *rpsL*. Drug resistance might thus be attributed to mutations located on another genomic region, such as *gidB* (51).

Several isolates harbouring high-confidence mutations were found to be phenotypically susceptible to the respective antibiotics. MiSeq and MinION sequencing analyses were repeated, and the same genotypic results were obtained. It should be noted that phenotypic susceptibility tests fundamentally measure the growth rate as a proxy for resistance. Slow growth (>21 days) during the DST may explain the false susceptibility in samples harbouring these mutations (52).

In this study, MinION sequencing required 6 hours, resulting in a total operation time of 15 hours. This was much shorter than the 38 hours required to complete the MiSeq workflow. A full-panel genotypic DST could be delivered for treatment guidance at least 1 day earlier than the MiSeq results, and approximately 18 days earlier than the pDST results.

A batch run of 12 samples on the MinION platform yielded a cost per sample of US$71.56, which excluded instrument and personnel costs. As discussed above, the batch size could be increased to a maximum of 24 samples, which would reduce the per-sample cost to a minimum of US$35.78. However, this price is not comparable to that of the MiSeq workflow, for which the high capacity for sample batching has driven down the costs. Specifically, the cost per sample in a MiSeq run was US$67.83 at a batch size of 24 samples. This cost could be reduced to approximately US$15 at a maximum batch size of 108 samples when using the MiSeq Reagent Nano Kit. Additionally, standard and micro versions of the MiSeq Reagent Kits are available. These versions have higher data outputs and enable larger batch sizes, both of which could eventually reduce the cost per sample even further.

However, we note that the numbers of samples in regional clinical centres that provide individualised care are unlikely to maximise the batching capacity of the MiSeq workflow. Instead, smaller batch sizes might decrease the time-to-results by reducing the time needed to collect a minimum number of samples per batch. Therefore, the reduced capital cost, simpler workflow and lower-throughput capabilities favour the MinION over the MiSeq in regional clinical settings that provide individualised care. Conversely, MiSeq is a better choice in high-throughput settings such as reference laboratories, where sample batching can be optimised to minimise costs at the expense of workflow complexity and time.

This study had several shortcomings. First, cultures of archived *M. tuberculosis* strains were used as the input samples for sequencing. However, the ability to conduct sequencing on patient samples directly (i.e., without requiring a culture) will be more appropriate for both patient care and surveillance. Second, phenotypic drug resistance profiles were not equally available for all isolates included in this study, and test results were unavailable for BDQ and LZD. This variable availability of data limited the evaluation of our sequence-based assay, particularly for these two drugs. Finally, although we compiled several established drug resistance-associated mutations in our drug resistance mutation panel, not all these mutations were present in our collection of clinical isolates. Therefore, we could not assess the ability of our assays to detect these rare mutations.

## Conclusions

Our study presented two targeted-sequencing workflows based on the Illumina MiSeq and nanopore MinION platforms for the rapid and comprehensive determination of drug-resistance in *M. tuberculosis* cultures. The diagnostic performance of the MinION platform was comparable to that of the commonly used MiSeq platform, which demonstrates the potential interchangeability of different sequencers in a laboratory setting. Both workflows enabled us to provide accurate and actionable results for the treatment of TB. Batching-based price constraints led to a higher sequencing cost per sample on the MinION platform than on the MiSeq platform under conditions of optimal batching. However, the MinION enabled a more open-access workflow, as fewer samples were batched per sequencing run. Therefore, we recommend the MiSeq workflow for high-throughput settings such as reference laboratories, and the MinION workflow for use in clinical settings that provide individualised care.

## Ethical Approval

The study protocols were approved by the ethical approval committees of the College of Health Arsi University (A/U/H/C/87/4016) and The Hong Kong Polytechnic University (HSEARS20161212001).

## Consent for publication

Not applicable

## Availability of data and materials

All raw sequencing data generated from MiSeq and MinION have been deposited under NCBI BioProject Sequence Read Archive (project accession PRJNA557083).

## Authors’ contributions

KT, TN, WY, GS conceived and designed the study; KT, TN, HL, RR, CC and GS performed laboratory experiments and sequencing; KL, KGT, BN, GA, and WY contributed *M. tuberculosis* cultures or phenotypic DST data; DG and CS developed the analysis pipeline; KT, TL, ST, LH and GS performed sequencing and statistical analysis; OM, WY and GK secured research funding for this project; KT, TN, HL and GS wrote/drafted and finalised the manuscript with contributions from all other authors. The final manuscript was read and approved by all authors.

## Acknowledgements

We wish to thank Oxford Nanopore Technologies Ltd. for their professional technical support and advices for nanopore sequencing and subsequent data analysis.

